# Surgical Removal of Visceral Adipose Tissue has Therapeutic Benefit in Male APP^NL-F^ Mice

**DOI:** 10.1101/2025.08.07.669175

**Authors:** Samuel A. McFadden, Yimin Fang, Kathleen Quinn, Mackenzie R. Peck, Jenelle E. Chapman, Tiarra Hill, Andrzej Bartke, Erin R. Hascup, Kevin N. Hascup

## Abstract

**Purpose:** Visceral white adipose tissue (vWAT) accumulation causes systemic inflammation, insulin resistance, metabolic syndrome, and senescent cell accumulation that are risk factors for Alzheimer’s disease (AD). Visceral fat removal (VFR) improves metabolism and reduces pro-inflammatory cytokines. We hypothesized that VFR removal in AD mice would improve metabolism and cognition.

**Methods:** Male and female APP^NL-F^ mice underwent sham or vWAT surgical resection (periovarian or epididymal and perirenal) at 4 (pre-symptomatic) and 16 (symptomatic) months of age to understand interventional and therapeutic effects, respectively. At 18 months of age, glucose metabolism and novel object recognition (NOR) memory were assayed followed by assessment of body composition and tissue-specific markers of metabolism, cell senescence, inflammation, or amyloid accumulation.

**Results:** Male and female APP^NL-F^ mice showed distinct VFR responses. In pre-symptomatic males, increased vWAT lipolysis and hepatic lipogenesis led to ectopic liver lipid accumulation, with reduced adiponectin and leptin, elevated visfatin, and impaired glucose metabolism. Symptomatic males showed reduced vWAT lipogenesis, enhanced hepatic lipolysis, glycolysis, and glycogenesis, lowering liver lipids and improving insulin sensitivity. Only symptomatic males improved NOR, linked to elevated hippocampal learning and memory markers. Female vWAT reaccumulation was due to increased lipogenesis and lower lipolysis. Pre-symptomatic females had lower hepatic lipogenesis, while glycolysis and glycogenesis declined with disease progression. Hippocampal senescence and inflammation were elevated early in the disease that persisted symptomatically.

**Conclusions:** Sex-specific differences in glucose and lipid metabolism and lipid accumulation underlie the divergent responses to VFR in APP^NL-F^ mice, with symptomatic males showing the only beneficial outcomes in metabolism and cognition.

**Graphical Abstract:** 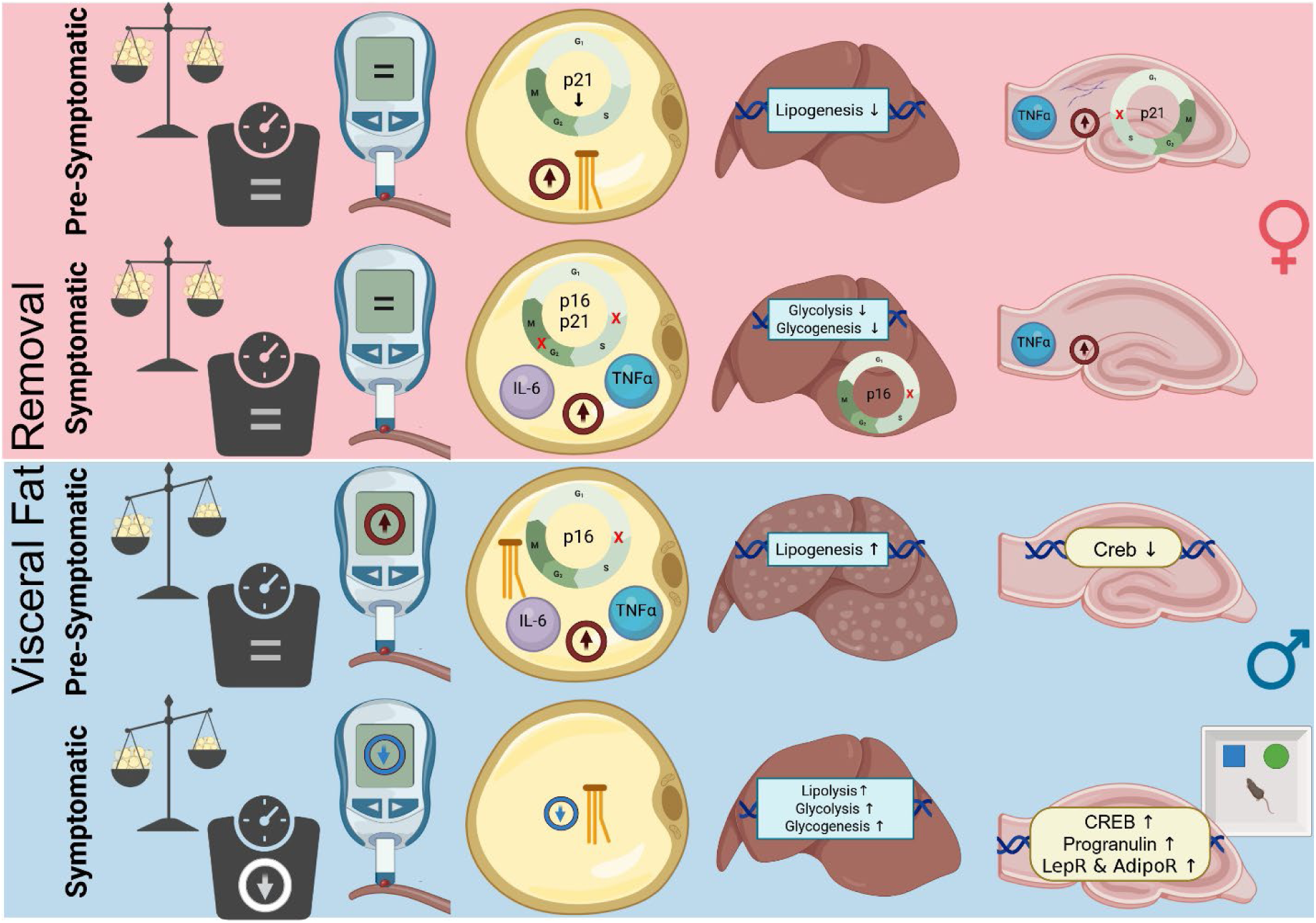

Male and female APP^NL-F^ mice exhibited distinct VFR responses to AD progression. In males, early disease stages were marked by vWAT lipolysis and hepatic lipid accumulation with metabolic dysfunction, while symptomatic stages showed a metabolic shift that improved insulin sensitivity and NOR performance. In contrast, females displayed progressive vWAT reaccumulation, reduced hepatic metabolism, and persistent hippocampal senescence and inflammation from early stages onward.

## Introduction

AD is a progressive, neurodegenerative disorder, and the most prevalent form of dementia. AD is historically characterized by senile plaques composed of aggregated amyloid beta (Aβ) and neurofibrillary tangles formed from hyperphosphorylated tau protein. However, AD is a multifactorial disorder characterized not only by aberrant protein aggregation, but also by mitochondrial dysfunction, oxidative stress, cellular senescence, and chronic inflammation - all of which contribute to disease progression (Breijyeh and Karaman, 2020). The deleterious cascade underlying AD progression is hypothesized to begin in midlife and is not isolated to the CNS. Peripheral organs, such as vWAT, regulate brain function through hormone secretion as well as glucose and lipid metabolism (Gustafson, 2012, Dove et al., 2021). Accumulation of vWAT in midlife is a predictor for future development of AD (Huang et al., 2022).

Adipose tissue regulates many aspects of whole-body physiology, including food intake, maintenance of energy levels, insulin sensitivity, body temperature, and immune responses (Sakers et al., 2022). Adipose tissue is located under the skin (subcutaneous; SubQ), between and around organs (vWAT), interscapular brown adipose tissue (BAT), in bone marrow, and in mammary glands (Wong Zhang et al., 2023). Beyond total adipose tissue mass, the distribution of fat depots is a stronger predictor of obesity-related disease risk and associated complications (Chusyd et al., 2016). Excess vWAT accumulation causes systemic inflammation, insulin resistance, reduced cerebral blood flow, and metabolic syndrome, which contributes to brain volume shrinkage and cognitive decline (Richard et al., 2000, Wong Zhang et al., 2023, Baglietto-Vargas et al., 2016, Biessels and Despa, 2018, Dove et al., 2021, Nguyen et al., 2014).

The link between systemic inflammation with vWAT and metabolic dysfunction has been reported in diabetic fatty rodents and in aged rats (Borst et al., 2005, Gabriely et al., 2002, Pitombo et al., 2006). In humans, vWAT is a stronger predictor of age-related cognitive impairment than body mass index (Whitmer et al., 2008, Elias et al., 2005, Kamogawa et al., 2010). A cross-sectional study also showed that vWAT dysfunction and inflammatory status in obesity, is associated with increased cerebral Aβ burden in midlife (Kim et al., 2022). Transplantation of vWAT from obese mice led to impaired cognition in lean wild-type recipient mice mediated by NLRP3/IL-1β signaling (Guo et al., 2020). The effects of VFR are broadly similar to observations in calorie restricted rodents (Masternak et al, 2012). VFR prevented obesity-induced insulin resistance, hyperinsulinemia, and decreased plasma pro-inflammatory cytokine levels (Borst et al., 2005, Gabriely et al., 2002, Pitombo et al., 2006), which improved glucose metabolism (Barzilai et al., 1999, Foster et al., 2011, Franczyk et al., 2021). Aging also increases vWAT senescent cell burden leading to a chronic pro-inflammatory state (Tchkonia et al., 2010). Reduction of senescent cells in vWAT improves metabolic and cognitive performance in normal aging (Fang et al., 2023) and AD mice (Fang et al, 2025).

Considering the contribution of vWAT accumulation on cognitive decline and AD progression, we hypothesized that surgical resection of this adipose depot would improve the metabolic profile and reduce cognitive impairments in the APP^NL-F^ AD mouse model. APP^NL-F^ mice contain the human Aβ isoform of the Swedish (KM670/671NL) and Iberian (I716F) mutations knocked into the *App* locus leading to upregulation of Aβ_42_ without overexpression of APP (Saito et al., 2014, Benitez et al., 2021). Plaque accumulation begins at ∼6 months of age while cognitive impairments occur between 12-18 months of age. Our prior research indicates that both sexes of APP^NL-F^ mice have increased markers of senescent cell (p16^Ink4α^ and p21^Cip1^) and senescence-associated secretory phenotype (SASP; IL-6, TNFα, MIP1α) in vWAT by 4 months of age (Fang et al., 2025), which may account for their reduced glucose tolerance (Abi-Ghanem et al., 2025). Besides metabolic dysfunction, senescent adipocytes also contribute to chronic systemic inflammation which diminishes cognitive reserve, promotes the accumulation of misfolded proteins, and accelerates AD progression (Huang et al, 2022).

To test our hypothesis, we surgically resected vWAT (periovarian or epididymal and perirenal, hereafter perigonadal) from male and female APP^NL-F^ mice at either 4 or 16 months of age. These time points allowed us to examine the effects of VFR for pre-symptomatic intervention (4 months) or symptomatic therapy (16 months). All metabolic and cognitive assays were conducted at 18 months of age. Our results indicate that VFR effects were disease stage- and sex-dependent with metabolic and cognitive benefits observed in symptomatic male APP^NL-F^ mice.

## Material and Methods

### Chemicals

All chemicals were stored according to manufacturer recommendations unless stated otherwise. The Phosphovanilin reagent for lipid determination was prepared by using 6 milligrams of vanillin (Fisher Scientific Cat # AC140821000) dissolved in 100 mL of 37°C H_2_O and further diluted to 500 mL with 85% phosphoric acid.

### Animals

Founder APP^NL-F^ mice on a C57BL/6 background (RRID: IMSR_RBRC06343) were obtained from Riken (Japan). C57BL/6 (RRID:IMSR_JAX:000664) mice were originally obtained from The Jackson Laboratory (Bar Harbor, ME) and used to maintain the genetic background of the breeding colonies. At weaning, pups were randomly assigned to mixed-litter cages and group-housed with mice of the same sex and genotype to control for litter size, maternal care, and other potential litter-specific variables. Protocols for animal use were approved by the Institutional Animal Care and Use Committee at Southern Illinois University School of Medicine (Protocol number: 2025-055), which is accredited by the Association for Assessment and Accreditation of Laboratory Animal Care. All studies were conducted in accordance with the United States Public Health Service’s Policy on Humane Care and Use of Laboratory Animals. Mice were group housed on a 12:12 h light-dark cycle, with food (Chow 5001 with 23.4% protein, 5% fat, and 5.8% crude fiber; LabDiet PMI Feeds) and water available *ad libitum*. Genotypes were confirmed by collecting a 5 mm tail tip for DNA analysis by TransnetYX, Inc (Cordova, TN). The number of mice in each cohort used for the study are shown in Table 1.

**Table 1:**
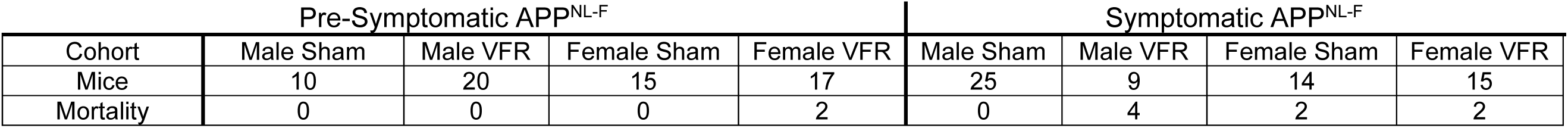
The number of mice in each cohort as well as the number of mice that experienced post-surgical mortality for that cohort.

### VFR Surgery

4 and 16-month-old male and female APP^NL-F^ mice were randomized into two groups – VFR (perigonadal) and sham-operated. Mice were isoflurane anesthetized, abdominal fur shaved, and betadine scrubbed over the incision site. An ∼1.5 cm vertical midline incision along the lower abdomen in males or ventral incisions on either lateral abdominal side in females were made to access epidydimal adipose depots. The perinephric adipose depots were removed by flank incisions. Adipose depots were removed by blunt dissection. Mice that underwent the sham operation had their vWAT mobilized, but not removed. Absorbable sutures were used to close the body cavity while wound clips were used on the skin. Tylenol (1 mg/mL) was administered in the drinking water from one day pre- to 3 days post-surgery. Table 1 shows the number of mice that were euthanized two weeks or less post-surgery for each group.

### Glucose tolerance test (GTT) and insulin tolerance test (ITT)

Glucose metabolism was assessed via GTT and ITT when mice reached 18 months of age (14 or 2 months post VFR surgery) as previously described (Fang et al., 2023). Fifteen-hour-fasted mice underwent GTT by intraperitoneal (ip) injection with 2.0 g glucose per kg of body weight (BW). Blood glucose levels were measured at 0, 15, 30, 45, 60, and 120 min with a PRESTO glucometer (AgaMatrix). For ITT, mice were injected ip with 1 IU insulin glargine (Sanofi-Aventis) per kg of BW. Blood glucose levels were measured at 0, 15, 30, 45, 60, and 120 min.

### NOR task

Object recognition memory was assessed when mice reached ∼19 months of age (15 or 3 months post VFR surgery). The NOR was used to evaluate memory retention based on the premise that mice spend more time exploring a novel object rather than a familiar object if the animal remembers the familiar object. Mice were habituated in the open field chamber for 30 min on the first day. 24 hours after, the mouse was returned to the chamber and presented with two similar objects for 5 min. A 24 hr inter-session-interval was used between introduction and retention phases to promote long-term memory retrieval. During the retention phase one of the familiar objects was replaced with a novel object and the mouse was given 5 minutes of exploration. The ANY-maze video tracking system (Stoelting Co., Wood Dale, IL) was used to record mouse navigation during the familiarization and retention phases and used to calculate the retention index and novelty preference.

### Body composition and tissue procurement

At 19 months of age, BW was recorded and unfasted mice were anesthetized using isoflurane. A cardiac puncture was used to collect the plasma. The blood was mixed with EDTA, followed by centrifugation at 10,000 g for 15 min at 4°C for plasma collection. Immediately following cardiac puncture, mice were decapitated and the peripheral tissues (liver, pancreas, perigonadal and SubQ, WAT, and BAT were weighed to calculate their contribution to total BW. Tissue was immediately flash frozen and stored at −80°C until processing as previously described (Fang et al., 2023).

### vWAT Histomorphology and Quantification

Frozen vWAT was transferred to ice cold 4% paraformaldehyde in phosphate buffer (pH 7.2) and fixed at 4°C for 72 hrs. Tissue was then cryoprotected in 30% sucrose in phosphate buffer (pH 7.2) and stored at 4°C for a minimum of 24 hrs. Following fixation, samples were embedded in optimal cutting temperature compound (Fisher Scientific Cat #23-730-571) using 15 mm cryomolds (Fisher Scientific Cat #22-363-553) on dry ice. Sections (10 µm) were cut using a Leica CM1950 cryostat (chamber −30°C; specimen holder −40°C) and mounted on gelatin-coated, charged glass slides (Fisher Scientific, Cat #12-550-15). Slides were air-dried at room temperature (∼25°C) for at least 15 minutes and stored at 4°C until staining. Slide-mounted vWAT sections were stained according to the manufacturer’s instructions using a commercial hematoxylin and eosin (H&E) staining kit (Abcam Cat # ab245880). Following staining and dehydration, sections were cleared with xylene substitute (Fisher Scientific Cat #9990505) and coverslipped with Permount mounting medium (Fisher Scientific Cat # SP15-100). Brightfield images were acquired at 40x magnification using a BZ-X810 All-in-One Flourescence microscope (Keyence Corp). For each sample, four fields of view from four separate section were imaged (16 images per sample). Adipocyte number and interior area (µm^2^) were quantified for each field using the Hybrid Cell Count software (Keyence Corp.). Measurements were averaged across fields and sections to generate a single mean adipocyte count and sample area per mouse.

### Total Liver Lipid Quantification

A modified micro-scale colorimetric sulfo-phospho-vanillin method was used to measure total liver lipids (Tran et al., 2019). Approximately 50 mg of liver tissue was homogenized in 1000 μL of chloroform/methanol (2:1) followed by addition of 500 μL of saline. The mixture was vortexed then centrifuged at 10,000 rpm for 5 minutes. The chloroform layer was separated and evaporated at 90°C for ∼20 min, then incubated at 90°C with 200 μL of 98% sulfuric acid for 20 min. After adding 750 μL of vanillin–phosphoric acid reagent, samples were incubated at 37°C for 15 min, cooled to 25°C, and 200 μL was transferred to a 96-well microplate. Absorbance was measured at 530 nm, and lipid content was normalized to tissue weight.

### Assessment of blood chemistry

Per the manufacturer’s protocol, adiponectin or leptin was measured with respective ELISA kits from Crystal Chem (Elk Grove Village, IL; Cat# 80569, 90030). Visfatin was measured by an ELISA kit from Mybiosource (San Diego, CA; Cat# MBS760980).

### RT–PCR

mRNA expression was analyzed by quantitative RT–PCR as previously described (Fang et al., 2020) using Bio-Rad cDNA synthesis kits (Cat # 1708897 and 1708891) and SYBR green mix (Cat # 1725121) and the QuantStudio 3 Real-Time PCR (Thermo Fisher Scientific). RNA was extracted using a RNeasy mini kit or RNeasy Lipid Tissue Mini Kit (Qiagen) following the manufacturer’s instructions. Relative expression was calculated as previously described (Masternak et al., 2012) and primers were purchased from Integrated DNA Technologies. A list of forward and reverse primers used in this study is shown in Table 2.

**Table 2:**
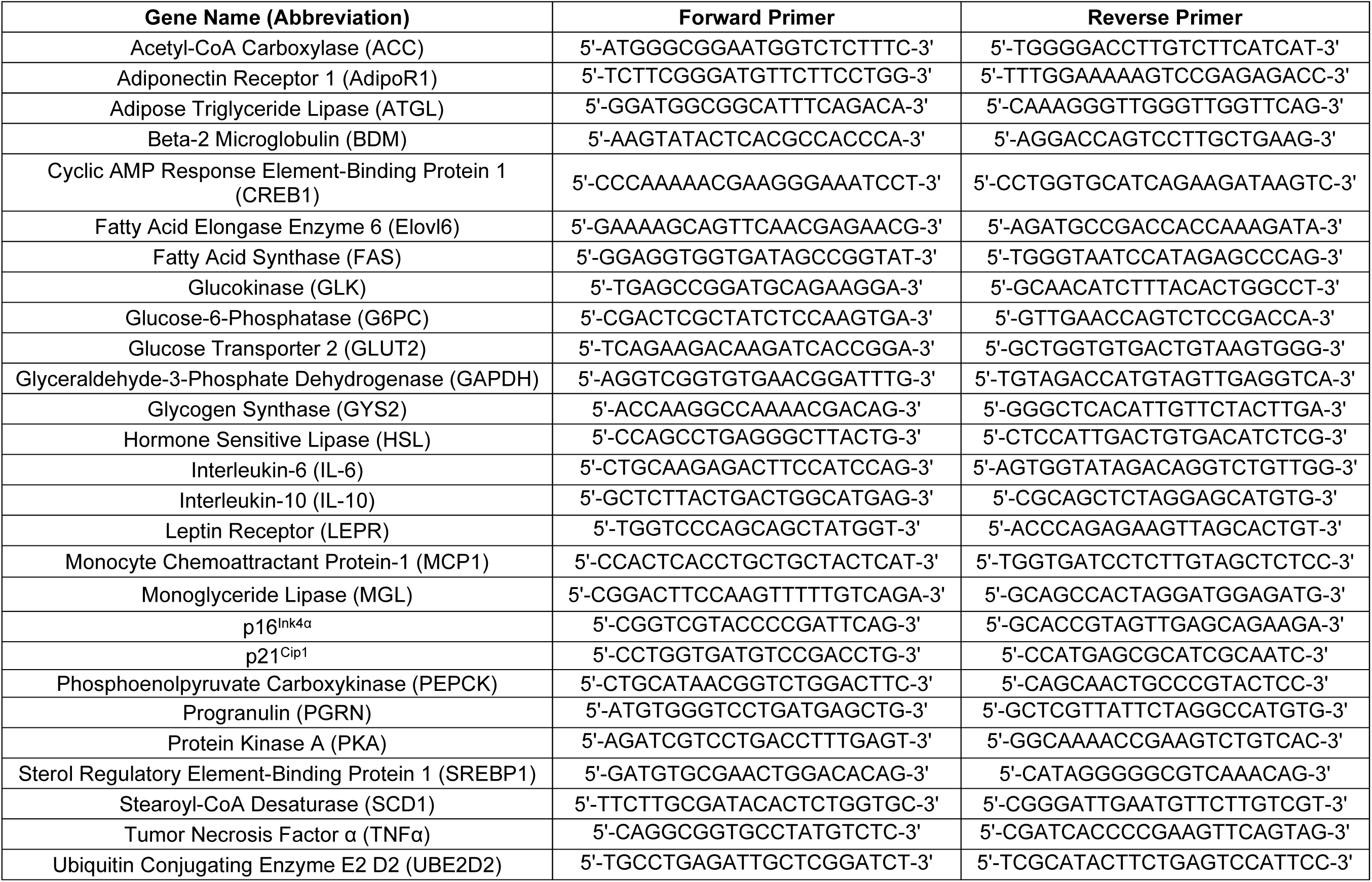
A list of forward and reverse primers.

### Soluble Aβ_42_ Quantification

The hippocampus from one hemisphere was dissected and stored at −80°C until tissue processing. Soluble Aβ_42_ concentrations were determined using the Human / Rat β amyloid ELISA kits (WAKO Chemicals; Cat: 290-62601) according to the manufacturer recommended protocols.

### Statistical analysis

Statistical analyses were conducted to determine differences between age- and sex-matched sham and VFR groups. Statistical tests and post hoc analyses are indicated in each Figure legend. A Grubb’s test was used to identify outliers. Significance was defined as p < 0.05 and investigatory as p < 0.10. Data are presented as mean ± SEM. All statistical analyses and graphs were completed using Prism 10 (GraphPad Inc, La Jolla, CA, USA).

## Results

### VFR at the symptomatic time point improved object recognition memory in male APP^NL-F^ mice

To explore whether VFR could improve object recognition memory in APP^NL-F^ mice, surgery was performed at a pre-symptomatic stage (4 months of age, prior to detectable plaque accumulation and cognitive decline) or a symptomatic stage (16 months of age, during advanced pathology and functional decline). Both male and female APP^NL-F^ cohorts were included to assess sex-specific effects of VFR. Object recognition memory was assayed at 18 months of age via the NOR task. Only male APP^NL-F^ mice that underwent VFR at the symptomatic time point had improved recognition memory as indicated by increased retention index (Fig. 1A) and novelty preference (Fig. 1B) compared to the age- and sex-matched sham surgery controls. VFR surgery performed at the pre-symptomatic time point in male, or either time point of female APP^NL-F^, had no effect on object recognition memory (Fig 1A-B).

**Figure 1.**
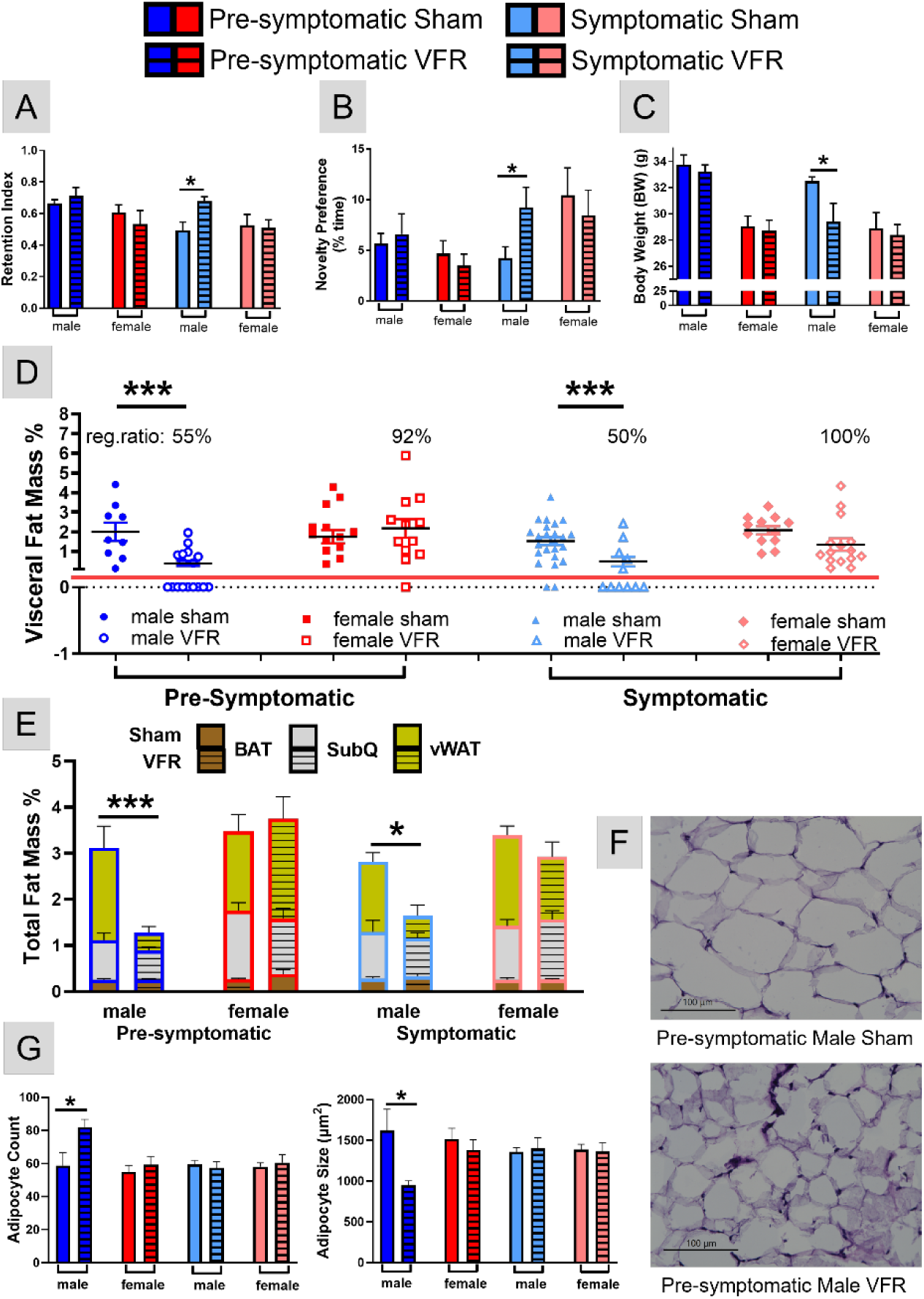
NOR and body composition analysis after VFR. The retention index (A) and novelty preference (B) for the NOR task were calculated at 18 months of age for all groups. BW (C) and the percentage of vWAT (D) and total fat mass for BAT, SubQ, and vWAT (E) in relation to BW were determined upon euthanization at 19 months of age. The percentage of VFR mice that regenerated vWAT (reg. ratio) are shown in (D). Representative 40x images of H&E stained vWAT in pre-symptomatic male sham (top) and VFR (bottom) cohorts (F). The average number of vWAT adipocytes and their internal diameter were determined for all groups (G). Data are represented as means ± SEM. *p<0.05, **p<0.01 based on a two-tailed Welch’s *t* test (A-C, E), a two-way ANOVA with Sidak’s post hoc (D), and an unpaired two-tailed t test (G); n=4-20.

### VFR resulted in changes to BW and composition in an age- and sex-dependent manner

To assess physiological changes caused by VFR, we examined body composition at ∼19 months of age upon euthanasia. VFR decreased BW at the symptomatic stage in male APP^NL-F^ mice compared to the sham group (Fig 1C). No differences were observed in the other VFR groups. The body composition analyses indicated that after VFR, vWAT accumulated in all ages and sexes but at different percentages. After VFR, APP^NL-F^ mice had vWAT reaccumulation occurring in 92% and 100% of mice in the pre-symptomatic and symptomatic cohorts, respectively (Fig. 1D). In contrast, ∼50% of the male mice undergoing VFR surgery at either age observed vWAT reaccumulation (Fig. 1D). This difference in vWAT was the major contributor to the observed changes of total percent adipose mass when including SubQ, interscapular BAT, and vWAT (Fig. 1E). In summary, the BW of symptomatic male APP^NL-F^ mice after VFR was reduced mainly due to less vWAT reaccumulation.

### Reaccumulated vWAT in pre-symptomatic male APP^NL-F^ mice had increased adipocyte number and smaller size

H&E staining was used to examine if reaccumulated vWAT had morphological differences. Representative 40x images of pre-symptomatic male sham and VFR APP^NL-F^ cohorts are shown in Fig. 1F. Reaccumulated vWAT following VFR in this cohort exhibited adipocyte hyperplasia characterized by a reduced mean internal diameter (Fig. 1G), and may account for the observed decrease in vWAT mass (Fig. 1E). No differences were observed in the other VFR treated groups compared to sham littermates.

### VFR during the pre-symptomatic time point in male APP^NL-F^ mice caused ectopic liver lipid deposition via increased de novo lipogenesis

Although vWAT reaccumulated similarly after VFR regardless of disease stage in male APP^NL-F^ mice, BW at the time of euthanasia was comparable between pre-symptomatic males after VFR and their sham controls. Body composition changes without effects on BW are known to happen with aging (St-Onge and Gallagher, 2010). We sought to identify tissue composition changes that caused the increased BW despite reduced adiposity in the pre-symptomatic male mice after VFR. Lipodystrophy leads to the excessive deposition of lipids in other organs, such as the liver or muscle, contributing to insulin resistance (Petersen and Shulman, 2018). To test whether the liver had more ectopic lipid deposition, the percentage of total liver lipids was measured. Pre-symptomatic male APP^NL-F^ mice had a higher percentage of liver lipids after VFR compared with sham controls (Fig 2A). In symptomatic APP^NL-F^ male mice, VFR reduced total liver lipids compared with sham surgery controls while no differences were observed in females at either disease stage.

**Figure 2.**
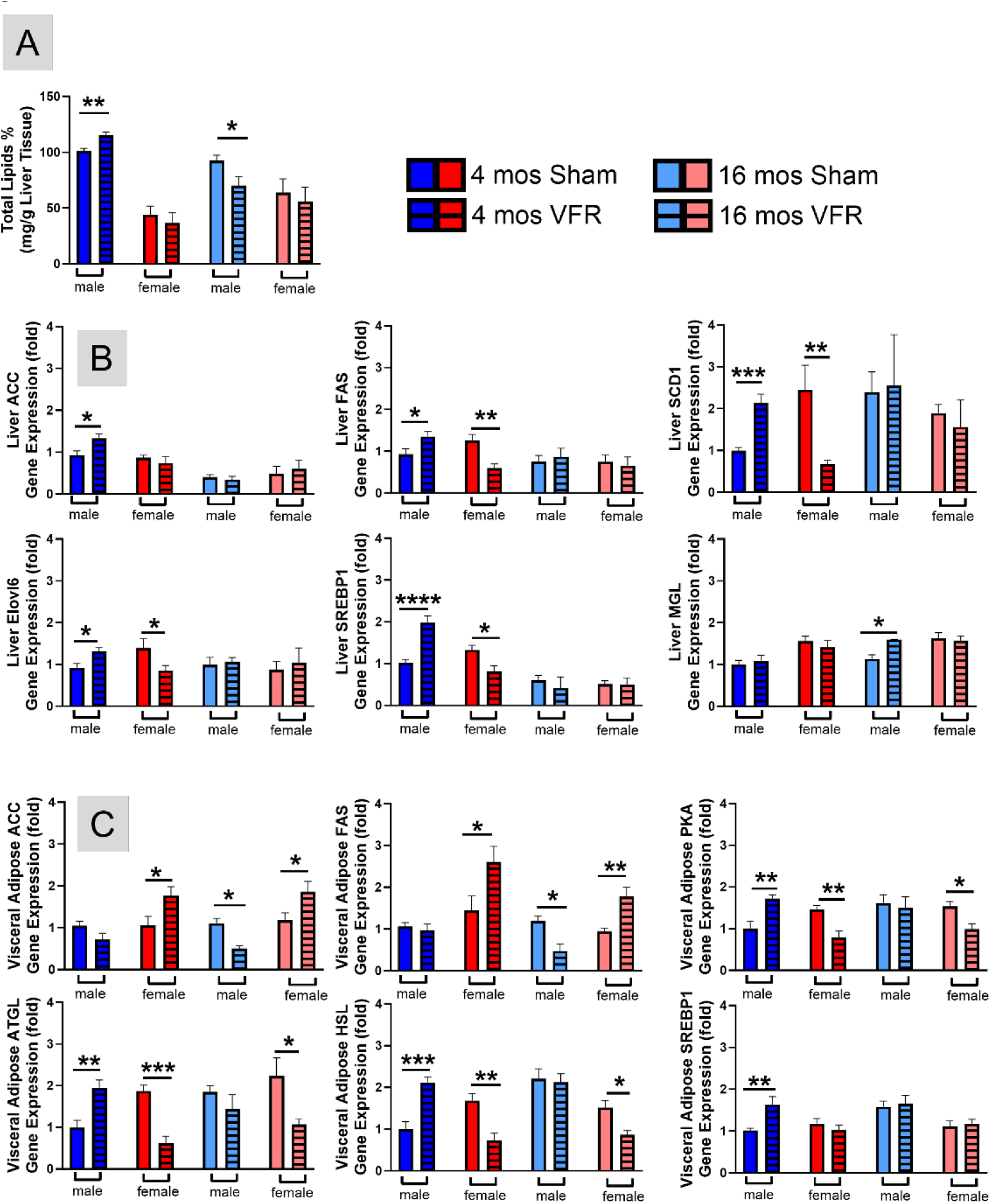
Lipid metabolism in hepatic and vWAT. The sulfo-phospho-vanillin method was used to measure the percentage of total liver lipids (A). qRT-PCR was used to determine relative gene expression profile of lipolysis and lipogenesis genes in the liver (B) and vWAT (C). Data are represented as means ± SEM. *p<0.05, **p<0.01, ***p<0.001, ****p<0.0001 based on a two-tailed Welch’s t test (n=8-15). Abbreviations: ACC – acetyl CoA-carboxylase, FAS – fatty acid synthase, SCD1 - stearoyl-CoA Desaturase, Elovl6 – fatty acid elongase enzyme 6, SREBP1 – sterol regulatory element-binding protein 1, MGL – monoglyceride lipase, PKA – protein kinase A, ATGL – adipose triglyceride lipase, HSL – hormone sensitive lipase

Liver gene expression showed that key *de novo* lipogenesis genes including acetyl-coenzyme A carboxylase (ACC), fatty acid synthase (FAS), stearoyl-CoA desaturase 1 (SCD1), elongation of long-chain fatty acids family member 6 (Elovl6), sterol regulatory element-binding protein-1 (SREBP1) were increased after VFR in pre-symptomatic male APP^NL-F^ mice compared with sham controls (Fig 2B). In pre-symptomatic female APP^NL-F^ mice, expression of these same genes (except for ACC) were decreased after VFR. Interestingly, liver gene expression of monoglyceride lipase (MGL), the rate-limiting enzyme in lipolysis, was only upregulated after VFR in symptomatic male APP^NL-F^ mice (Fig. 2B). This is one explanation for the decreased BW and vWAT observed in this group. These data indicated that VFR in pre-symptomatic male APP^NL-F^ mice cause more ectopic liver lipids likely due to higher *de novo* lipogenesis, while the contrary was observed in males at the symptomatic disease stage due to higher lipolysis.

### VFR in female APP^NL-F^ mice led to increased gene expression of de novo lipogenesis and decreased expression of lipolysis genes in reaccumulated vWAT

Nearly all the APP^NL-F^ female mice reaccumulated vWAT after VFR, with similar morphology to levels observed in sham surgery controls matched for age and sex (Fig. 1D, G). To understand the reason for this reaccumulation, we examined gene expression of lipogenesis and lipolysis in vWAT (Fig 2C). Expression levels of two key *de novo* lipogenesis genes, ACC and FAS, were increased in APP^NL-F^ female mice at either disease stage after VFR. The expression of ACC and FAS were decreased following VFR in male APP^NL-F^ mice at the symptomatic, but not at the pre-symptomatic time point. Lipolysis gene expression, including protein kinase A (PKA), adipose triglyceride lipase (ATGL), and hormone sensitive lipase (HSL), were decreased at both disease stages after VFR in APP^NL-F^ female mice compared with sham controls. In male mice, VFR increased the expression of PKA, ATGL, and HSL at the pre-symptomatic, but unchanged, at the symptomatic time point. The data suggests that the accumulation of vWAT after VFR in female APP^NL-F^ mice at either stage of disease progression is a result of increased lipogenesis and decreased lipolysis gene expression. In male APP^NL-F^ mice, the reduced accumulation of vWAT after VFR was disease stage specific with pre-symptomatic time points increasing expression of lipolysis genes, while surgery at the symptomatic time point decreased lipogenesis gene expression. Interestingly, SREBP1 vWAT expression was upregulated after VFR only in pre-symptomatic APP^NL-F^ male mice. SREBP1 is a transcription factor activated by high blood glucose (Bertolio et al., 2019, Zhao et al., 2024), suggesting altered glucose metabolism in this cohort.

### VFR surgery impacted either glucose tolerance or insulin sensitivity in male APP^NL-F^ mice

Lipid metabolism plays an important role in regulating glucose utilization, and the two processes are tightly interconnected (Parhofer, 2015). Accordingly, the observed changes in lipid metabolizing gene expression could affect glucose metabolism. To test the effects of VFR on blood glucose metabolism, we monitored fasting and fed glucose levels as well as glucose tolerance with a GTT, and insulin sensitivity with an ITT, at 18 months of age. VFR during the pre-symptomatic disease stage in APP^NL-F^ male mice reduced their glucose tolerance, but had no effects in symptomatic males (Fig 3A). VFR improved insulin sensitivity in symptomatic APP^NL-F^ male mice, but effects were not observed during the pre-symptomatic stage in males (Fig 3B). Consistent with the GTT, VFR increased fasting glucose in pre-symptomatic APP^NL-F^ male mice compared with sham controls (Fig. 3C). VFR did not affect fasting blood glucose in symptomatic APP^NL-F^ male mice, nor were differences in fed glucose observed in either disease stage of the male cohorts (Fig. 3D). Blood glucose levels were unchanged after VFR in either disease stage of APP^NL-F^ female mice.

**Figure 3.**
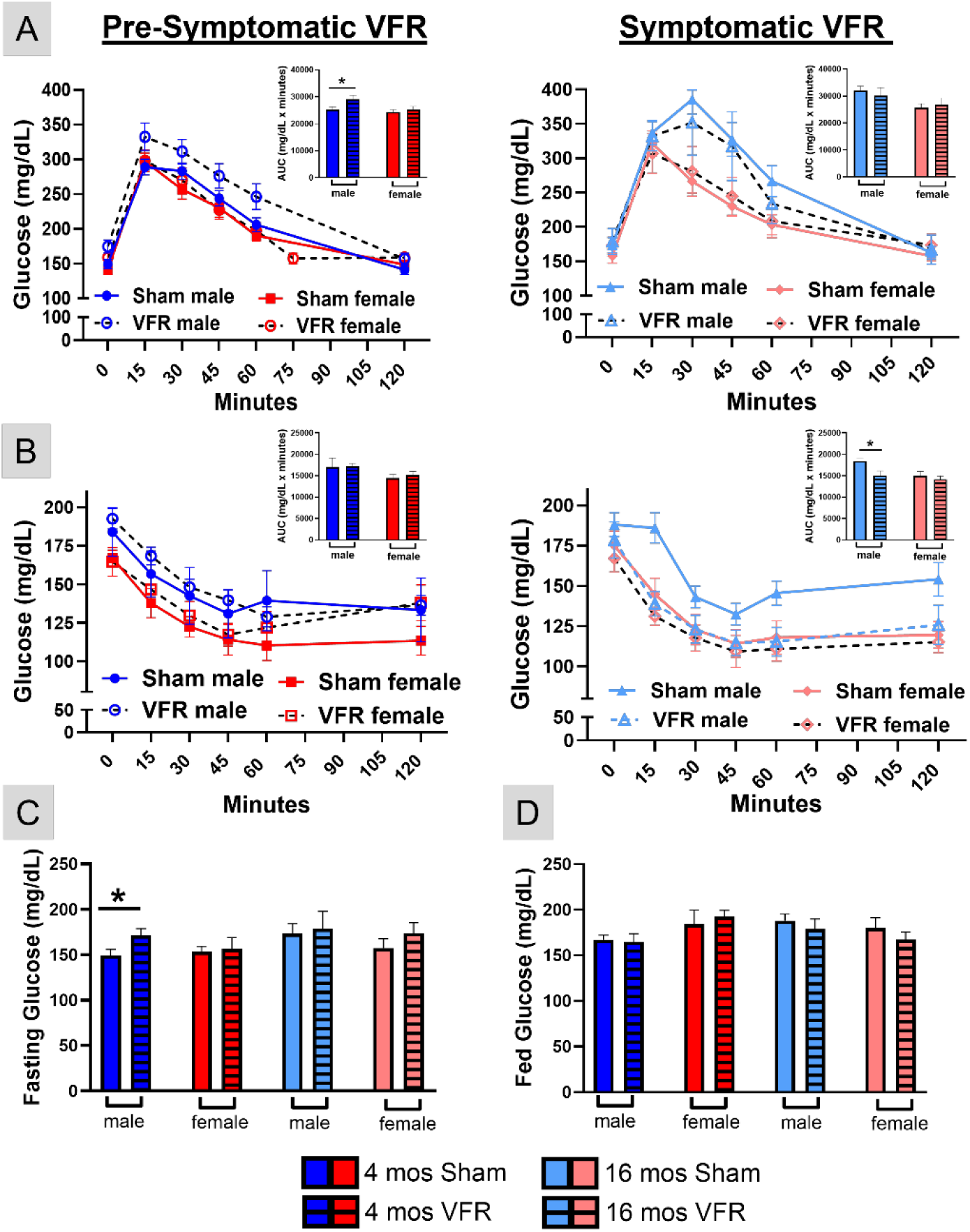
Blood glucose levels in pre-symptomatic and symptomatic APP^NL-F^ mice. Mice were fasted for 15 hours before they received an ip injection of glucose and blood glucose levels were monitored with a glucometer for two hours to determine glucose tolerance (A). Unfasted mice received an ip injection of insulin and blood glucose levels were monitored with a glucometer for 2 hours to determine insulin sensitivity (B). The area under the curve (AUC) for each assay are inset in pre-symptomatic (left) and symptomatic (right) cohorts. Fasting (C) and fed (D) blood glucose levels were determined from t=0 during the GTT and ITT, respectively. Data are represented as means ± SEM. *p<0.05 based on a two-way ANOVA with Sidak’s post hoc (A-D) and a two-tailed Welch’s t test (n=9-16).

The liver plays a major role in the control of whole body glucose homeostasis (Han et al., 2016, Guerra and Gastaldelli, 2020, Scoditti et al., 2024). To understand the observed blood glucose differences, we examined liver gene expression (Fig 4A) associated with gluconeogenesis (synthesis of glucose from non-carbohydrate sources), glycolysis (breakdown of glucose for energy), and glycogenesis (conversion of glucose to glycogen). VFR increased expression of gluconeogenesis genes, glucose-6-phosphatase (G6PC) and phosphoenolpyruvate carboxykinase (PEPCK), in pre-symptomatic male mice, consistent with their increased fasting blood glucose levels (Fig 3C). In contrast, glucose transporter 2 (Glut2) was increased in symptomatic males, while GCK (Glucokinase) and glycogen synthase 2 (GYS2) key genes in glycolysis and glycogenesis, respectively, were decreased in female APP^NL-F^ mice after VFR.

**Figure 4.**
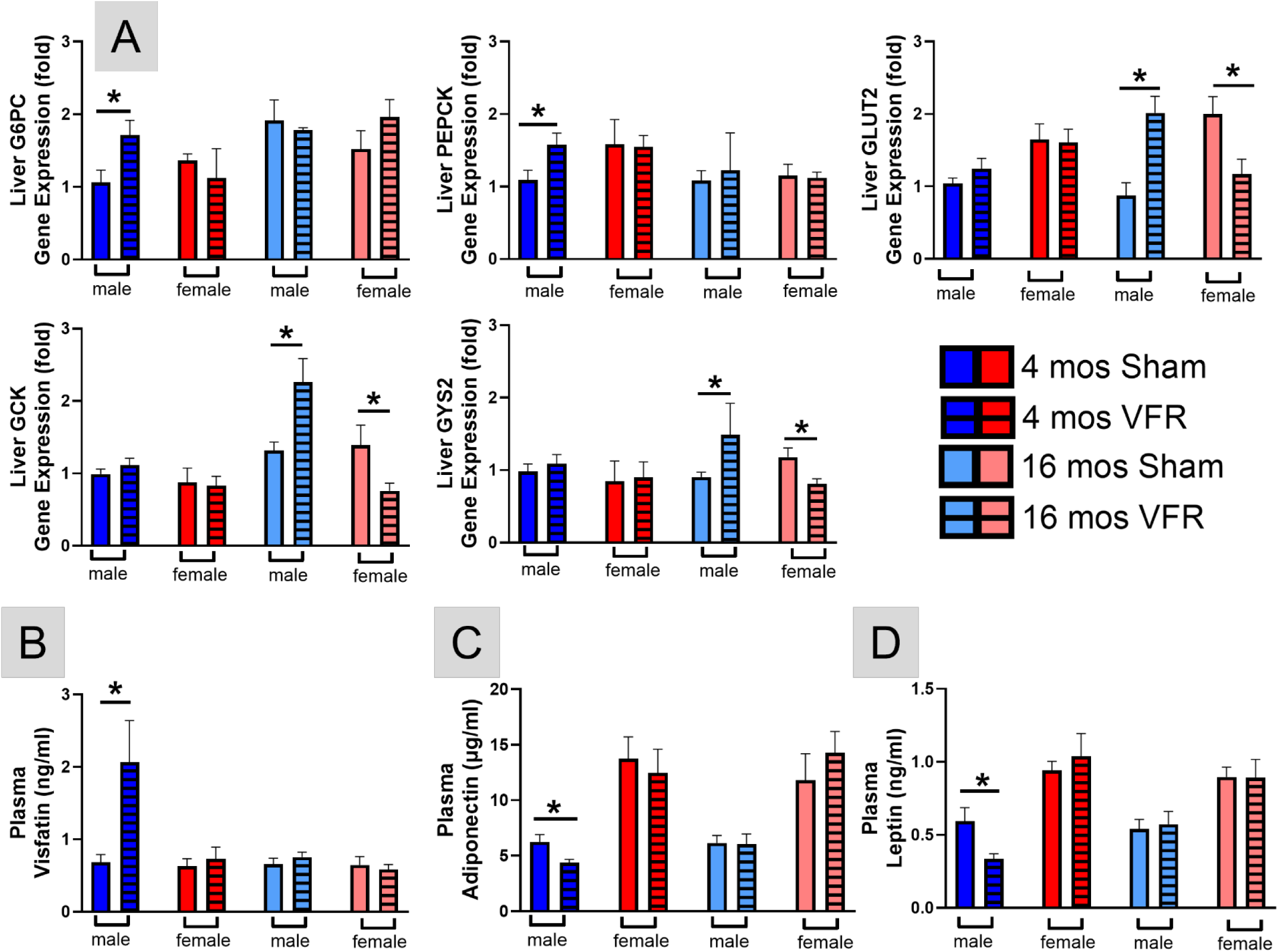
Liver glucose metabolism and plasma adipokines. qRT-PCR was used to determine hepatic relative gene expression profiles (A) of gluconeogenesis (G6PC, PEPCK), glycolysis (GLUT2, GCK), and glycogenesis (GYS2). Circulating visfatin (B), adiponectin (C), and leptin (D) determined by ELISA from plasma collected at the time of euthanasia. Data are represented as means ± SEM. *p<0.05 based on a two-tailed Welch’s t test (n=4-12). Abbreviations: G6PC - glucose-6-phosphatase, PEPCK - phosphoenolpyruvate carboxykinase, Glut2 - glucose transporter 2, GCK - glucokinase, GYS2 - glycogen synthase 2 (GYS2)

Taken together, the blood glucose and liver gene expression measures indicate VFR affected glucose metabolism in a sex- and disease stage-dependent manner. Pre-symptomatic APP^NL-F^ male mice had increased fasting blood glucose and made them less glucose tolerant after VFR, likely mediated via increasing gluconeogenesis. At symptomatic time points, VFR improved glucose metabolism by increasing insulin sensitivity via enhancing glycolytic and glycogenesis gene expression. Although this gene expression was reduced after VFR in symptomatic female mice, no effects on insulin sensitivity or blood glucose levels were observed.

### VFR alters plasma visfatin, adiponectin, and leptin in pre-symptomatic male APP^NL-F^ mice

Adipokines secreted from adipose tissue are important for whole body glucose metabolism and insulin sensitivity (Rabe et al., 2008, Kojta et al., 2020). Given the altered lipid and glucose metabolism after VFR in the APP^NL-F^ mice, we examined plasma adipokines. Visfatin is primarily produced in vWAT, with plasma levels increasing in response to high-fat diet feeding and obesity (Adeghate, 2008, Sethi and Vidal-Puig, 2005), and is associated with glucose intolerance (Abdalla, 2022). VFR increased plasma visfatin in pre-symptomatic male APP^NL-F^ mice compared with age-matched sham controls (Fig 4B), consistent with their higher ectopic liver accumulation, fasting blood glucose, and glucose intolerance. No plasma visfatin differences were observed in the other mouse cohorts.

Adiponectin enhances glucose and lipid metabolism (Yanai and Yoshida, 2019), decreases lipogenesis (Richard et al., 2000), and has neuroprotective effects (Khoramipour et al., 2021). VFR reduced plasma adiponectin in pre-symptomatic VFR APP^NL-F^ male mice (Fig 4C). This would also contribute to the reduced glucose metabolism and increased liver lipids observed in this cohort. Plasma adiponectin changes were not observed after VFR in the other cohorts.

Leptin regulates appetite and energy expenditure, and its deficiency can disrupt glucose and lipid metabolism, leading to neuronal damage that impairs neuroplasticity and memory consolidation (Fernandes et al., 2024). Higher leptin levels are also associated with a lower dementia risk (Lieb et al., 2009). Plasma leptin was decreased in pre-symptomatic APP^NL-F^ male mice after VFR compared with age-matched sham controls (Fig. 4D). This likely contributed to the imbalance of glucose and lipid metabolism in this male cohort. Differences in plasma leptin levels were not observed in the other cohorts of mice.

### VFR increases WAT senescent cell and SASP markers in pre-symptomatic male and symptomatic female APP^NL-F^ mice

Adipose tissue, particularly vWAT, has the highest burden of senescent cells (Palmer et al., 2021) and we previously showed senescent cell markers in vWAT were increased in male and female APP^NL-F^ mice (Fang et al., 2025). VFR can decrease systemic and neuroinflammation in aged mice (Shin et al., 2015). We examined gene expression of senescent cell and SASP markers in vWAT (Fig 5A). Gene expression of p16^Ink4α^, p21^Cip1^, and IL-10 were increased in vWAT after VFR in pre-symptomatic APP^NL-F^ male mice. Expression of vWAT p16^Ink4α^ and TNFα were also increased in symptomatic APP^NL-F^ female mice (Fig 5A) while no differences in IL-6 and MCP1 were observed in any cohort (data not shown) after VFR. SASP markers were also examined in SubQ WAT where VFR increased TNFα, IL-10, IL-6, and MCP1 in pre-symptomatic APP^NL-F^ male mice (Fig 5B). SubQ IL-6 was also increased in symptomatic APP^NL-F^ female mice after VFR. Overall, VFR increased gene expression of senescent cell and SASP markers in WAT of pre-symptomatic APP^NL-F^ male mice, with a modest increase observed in symptomatic females.

**Figure 5.**
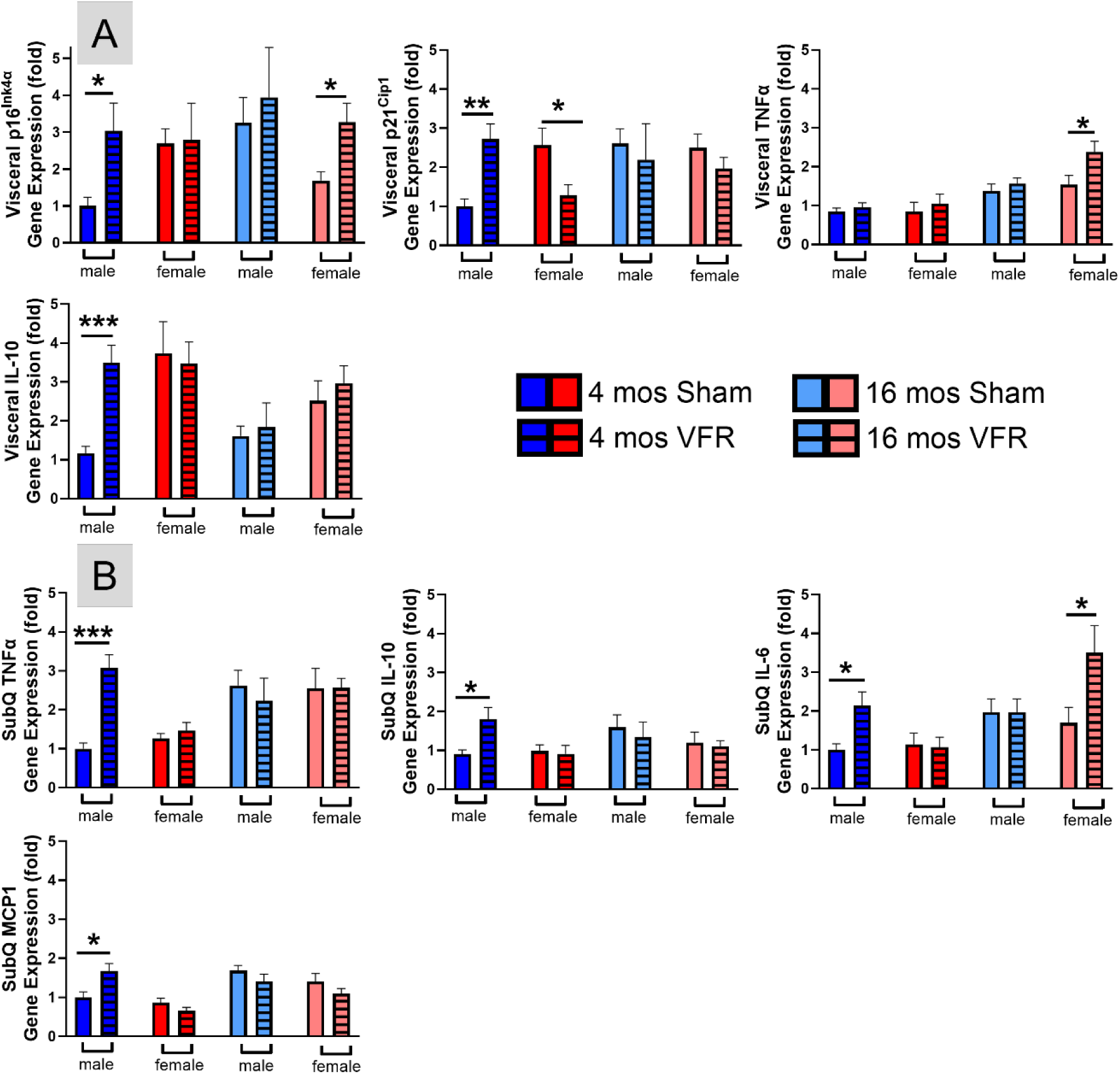
Senescent cell and inflammation markers in vWAT and SubQ. qRT-PCR was used to determine vWAT (A) and SubQ (B) relative gene expression profiles of senescence (p16^Ink4α^, p21^Cip1^) and inflammation (TNFα, IL-10, IL-6, MCP1). Data are represented as means ± SEM. *p<0.05, **p<0.01, ***p<0.001, ****p<0.0001 based on a two-tailed Welch’s t test (n=4-12). Abbreviations: TNFα – tumor necrosis factor alpha, IL – interleukin, MCP1 – monocyte chemoattractant protein 1.

### VFR modulates gene expression of hippocampal senescence, SASP, and cognitive markers

Excess vWAT accumulation is linked to accelerated brain aging, neuroinflammation, and dementia risk (Moser and Pike, 2016, Oh et al., 2023, Budamagunta et al., 2023). To understand how VFR affected hippocampal function, the brain region strongly associated with learning and memory, we examined gene expression of cell senescence and cytokines (Fig 6A). VFR had no effect on p16^Ink4α^, p21^Cip1^, or TNFα expression in APP^NL-F^ male mice at either stage of disease progression. Alternatively, p16^Ink4α^ gene expression was increased after VFR in pre-symptomatic APP^NL-F^ female mice. VFR caused a corresponding TNFα gene expression increase in this cohort as well as the symptomatic females. VFR had deleterious effects on hippocampal senescence and inflammation in female, but not male APP^NL-F^ mice.

**Figure 6.**
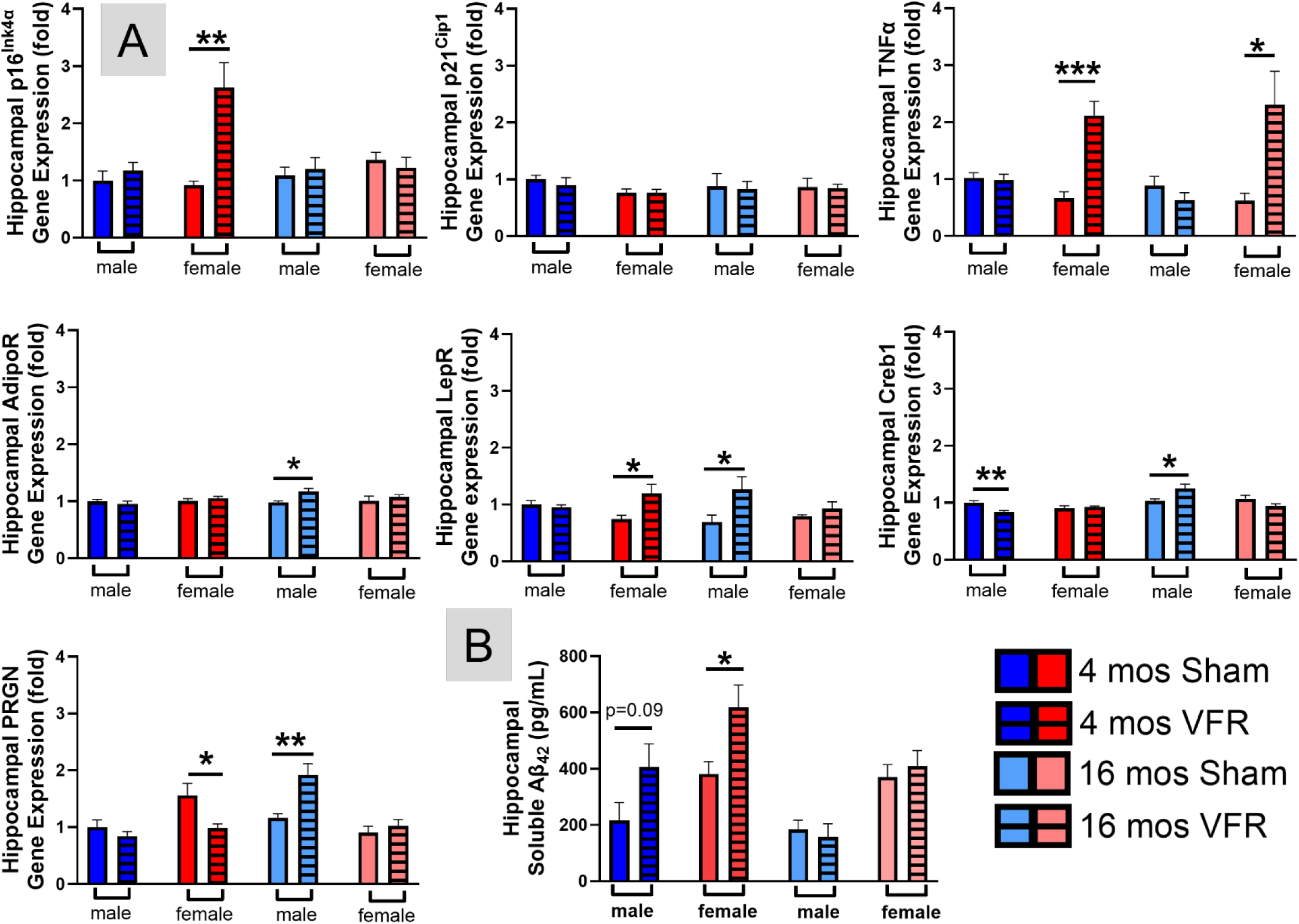
Senescent cell and inflammation markers in the hippocampus. qRT-PCR was used to determine hippocampal relative gene expression profiles (A) of senescence (p16^Ink4α^, p21^Cip1^), inflammation (TNFα), and memory (AdipoR, LepR, Creb1, PRGN), while soluble Aβ_42_ (B) was determined by ELISA. Data are represented as means ± SEM. *p<0.05, **p<0.01 based on a two-tailed Welch’s t test (n=4-12). Abbreviations: TNFα – tumor necrosis factor alpha, AdipoR – adiponectin receptor, LepR – leptin receptor, Creb1 – cyclic AMP Response Element-Binding Protein 1, PRGN – progranulin.

Since we observed improvements in object recognition memory after VFR in symptomatic male APP^NL-F^ mice, we examined markers associated with learning and memory in the hippocampus (Fig 6A). Both adiponectin and leptin receptors (AdipoR, LepR) are found in the hippocampus and play a role in synaptic plasticity and cognition. Gene expression of both receptors were upregulated after VFR in symptomatic APP^NL-F^ male mice (Fig 6A). In females, VFR increased LepR in the pre-symptomatic disease stage. Cyclic AMP response element-binding protein 1 (Creb1), plays a role in long-term memory formation. VFR decreased Creb1 gene expression in pre-symptomatic APP^NL-F^ male mice, but elevated expression in the symptomatic male cohort. Progranulin (PRGN) is important for neuronal survival and growth in the hippocampus. PRGN gene expression was decreased after VFR in pre-symptomatic APP^NL-F^ female mice, but elicited an increase in symptomatic males. Overall, VFR altered the hippocampal gene expression profile of symptomatic APP^NL-F^ male mice in a manner that would improve their object recognition memory.

### VFR increases soluble Aβ_42_ in pre-symptomatic male and female mice

Soluble Aβ_42_ is the neurotoxic species associated with synaptic dysfunction and cognitive decline in AD (Findley et al., 2019). An ELISA indicated hippocampal soluble Aβ_42_ (Fig 6B) production was only increased in pre-symptomatic stage of APP^NL-F^ male and female mice after VFR. VFR did not affect soluble Aβ_42_ during the symptomatic stage in either sex.

## Discussion

Adipose tissue accumulation and distribution exhibit sex-specific differences, yet these patterns are evolutionarily conserved across humans and mice (Chusyd et al., 2016). Females typically have a higher overall fat percentage with a large proportion stored subcutaneously for energy supply during pregnancy and lactation during food scarcity, while males accumulate fat in visceral regions. Numerous variables influence adipose development including sex chromosomes, gonadal hormone levels, and hormone receptors. Sex chromosome complement impacts adipose accumulation whereby the presence of two “X” chromosomes leads to higher accumulation while the presence of the “Y” chromosome has no influence. The gonadal hormones, testosterone and estrogen, drive vWAT and SubQ accumulation in males and females, respectively. Age-related hormonal changes increase vWAT accumulation regardless of sex and is a key factor in the development of metabolic syndrome that increases dementia risk.

Besides its role as an energy reservoir, adipose tissue is an endocrine organ that secretes peptides such as TNFα, IL-6, and adiponectin, which can influence either tissue-specific or whole body metabolism. Prior studies examining VFR effects in rodents were conducted to elucidate mechanisms contributing to metabolic improvements. VFR improves insulin sensitivity and glucose tolerance in young, aged, and obese rodents with some of the metabolic changes occurring through decreased free fatty acids, leptin, and TNFα expression (Kim et al., 1999, Gabriely et al., 2002, Shi et al., 2007a). VFR in aged C57BL/6 mice led to reduced expression of pro-inflammatory cytokines in the plasma, WAT, and cortex (Shin et al., 2015). Although adiponectin plays a key role in glucose and lipid metabolism and suppresses inflammatory cytokines, serum levels are found to decrease following VFR in male and female Sprague Dawley rats (Borst et al., 2005, Ling et al., 2014), which may be due to the absence of these WAT depots. Considering the metabolic benefits of VFR and the known link between metabolic dysfunction and AD risk, we aimed to investigate the impact of VFR in an AD model.

Surprisingly, the VFR response was disease stage- and sex-dependent. The removal of vWAT in symptomatic APP^NL-F^ male mice improved object recognition memory potentially due to increased hippocampal AdipoR, LepR, Creb1, and PRGN expression. Lipogenesis vWAT gene expression was down-regulated, while hepatic genes associated with lipolysis and glucose transport were up-regulated. These effects were associated with a decreased percentage of total liver lipids. All of these alterations were reflected by a reduced percentage of total fat, specifically reaccumulated vWAT, BW, and improved blood glucose maintenance. The opposite was true for VFR during the pre-symptomatic stage in males. After VFR, an imbalance of lipid metabolism was prevalent, reflected by an elevated expression of vWAT lipolysis and hepatic lipogenesis genes. This tissue metabolic imbalance may be a result of adipocyte hyperplasia with reduced cell size as well as the redistribution of lipids into other organs, such as the liver, where higher ectopic fat accumulation was observed. Plasma adiponectin and leptin were decreased along with increased visfatin resulting in impaired glucose metabolism and peripheral inflammation. Typically, adiponectin and leptin are inversely related, but plasma levels of both were decreased in pre-symptomatic male APP^NL-F^ mice. The decrease in leptin might be due to the loss of the adipose depot while the anticipated compensatory increase in adiponectin may not have been observed because the adiponectin source was also depleted. Further studies are needed to determine why this effect was not observed in the other cohorts after VFR. In the brain, reduced hippocampal Creb1 expression and increased soluble Aβ_42_, combined with the aforementioned adverse physiological changes, failed to improve object recognition memory.

In contrast to males, vWAT reaccumulated in nearly all female mice and was similar in number and size to sham littermates regardless of when VFR occurred. This could be attributed to higher lipogenesis and lower lipolysis gene expression. Although ectopic lipid deposition did not accumulate in the liver, VFR in pre-symptomatic female APP^NL-F^ mice led to decreased hepatic lipogenesis gene expression, while expression of genes involved in glycolysis and glycogenesis were decreased in symptomatic females. These results were in opposition to age-matched males. Hippocampal soluble Aβ_42_, senescent cell, and inflammatory markers were elevated in pre-symptomatic APP^NL-F^ female mice while the later was only observed at the symptomatic time point. No changes to blood glucose levels or object recognition memory were observed in females after VFR.

Lipectomy in mice, rats, and other small mammals does not result in long-term BW loss with regain occurring 12-14 weeks post-surgery (Murillo et al., 2019). In the majority of the studies this was attributed to regeneration of the lipectomized depot, redistribution of fat to other adipose depots, or ectopic fat accumulation in other organs. In our present study, female APP^NL-F^ mice regained BW due to reaccumulating vWAT. Although vWAT accumulation was lower in pre-symptomatic males, they regained BW due to ectopic fat distribution in the liver and possibly other organs. Only symptomatic APP^NL-F^ male mice had lower BW attributed to less visceral adiposity. Redistribution of fat to other adipose depots was not observed in any cohort. Our present study was designed to examine disease stage-specific interventional or therapeutic AD strategies rather than examining the contribution of aging on adiposity and AD progression. Regardless, effects of aging on adipose accumulation likely contributed to the observed differences in lipid distribution and the comparable end-of-study BW between sham and VFR cohorts matched for age and sex. The gradual decline in estrous cycling and the associated drop in estrogen levels, which promotes visceral rather than SubQ accumulation, may partially explain post-resection visceral accumulation in female mice, whereas aging males exhibit ectopic liver lipid deposition (Donato et al., 2014).

The exact mechanisms behind this adipose rebound are not well known, but suggest biological feedback systems that act to preserve energy balance. For example, the adipostat hypothesis postulates that alterations to energy reserves triggers compensatory mechanisms in metabolism and/or thermogenesis to restore the reservoir. Since gonadal hormones and sex chromosome complement mediate adipose storage, distribution, and utilization, male and female mice use different behavioral strategies to restore fat after lipectomy. Males increase caloric consumption while females reduce heat production and energy expenditure (Shi et al., 2007b). Our findings are in support of this whereby pre-symptomatic APP^NL-F^ male mice had increased gluconeogenesis gene expression and fasting blood glucose, and worse glucose tolerance after VFR. While VFR in Female APP^NL-F^ mice reduced glycolysis and lipolysis, but increased lipogenesis gene expression. It is important to note, however, that our study was not intended to serve as a preclinical model for liposuction, as that procedure primarily targets SubQ rather than vWAT depots. However, lipectomy in humans causes a transient decrease in BW and adipose depots that gradually return to preoperative levels within months (Hernandez et al., 2011, Seretis et al., 2015), suggesting the preservation of this energy balance is conserved across species.

The accumulation of WAT and subsequent increases in BW begins in midlife and continues into early aging in both humans (40-65 years) and mice (12-18 months). The reason for this increase is attributed to adipogenesis and adipocyte hypertrophy that are prominent during midlife (Wang et al., 2025). Remodeling of vWAT, like its distribution, varies between sexes, with estrogen levels playing a key role in regulating these processes and shifting notably with age. Female mice have increased adipose progenitor cells leading to hyperplasia while males experience hypertrophy (Wu et al., 2017). Both increase adipose accumulation, but hyperplasia is associated with better metabolic health (Steiner and Berry, 2022), which may explain why blood glucose levels were unaffected in female mice after VFR. Hypertrophic adipocytes secrete reduced adiponectin and increased pro-inflammatory cytokines, the latter promoting ectopic lipid distribution into peripheral organs. These pathological features were evident following VFR in pre-symptomatic APP^NL-F^ male mice, despite the reaccumulated vWAT exhibiting a less hypertrophic phenotype. Accordingly, adipose cell size rather than number correlates with increased risk for metabolic syndrome (Vishvanath and Gupta, 2019).

Senescent cell accumulation in both the periphery and brain contribute to aging and age-related neurodegenerative disorders such as AD. The mechanisms behind this are multifaceted and dependent upon the type of senescent cell (Zhu et al., 2024). Senescent brain cells alter amyloid precursor protein processing by shifting it from the non-amyloidogenic to the amyloidogenic pathway, promoting the formation of Aβ (Sun et al., 2018). The pro-inflammatory SASP response coupled with decreased microglial phagocytosis of misfolded proteins can lead to the subsequent Aβ accumulation and deposition in the brain. We observed elevated hippocampal soluble Aβ_42_ after VFR in pre-symptomatic APP^NL-F^ mice compared with sex-matched sham controls, which was contradictory to our hypothesis. VFR increased WAT senescent cell burden and SASP in pre-symptomatic males, while these were increased in the hippocampus of pre-symptomatic females. Considering senescent cell and SASP increases were predominantly observed in the pre-symptomatic groups, this may indicate a long-term effect of VFR. In AD mouse models, reducing peripheral and hippocampal senescent cell burden with senolytics has been shown to reduce amyloid pathology and improve cognitive performance - albeit with benefits favoring female mice (Currais et al., 2014, Zhang et al., 2019, Fang et al., 2025, Ng et al., 2024).

Given that AD pathology is thought to initiate in midlife prior to cognitive decline, the concurrent increase in adipose tissue accumulation during this period may play a causal role in disease progression. Our present study indicates accumulation or redistribution of adipose tissue during the pre-symptomatic AD stage can cause metabolic disturbances resulting in sex-specific peripheral and cerebral senescent cell accumulation that exacerbates pathological AD progression. However, during symptomatic AD stages, elimination of vWAT has beneficial metabolic and cognitive effects in male, but not female APP^NL-F^ mice. These sex-specific differences are attributed to increasing energy storage in female symptomatic mice.

Several study limitations should be noted when interpreting the results of the present study. Although the study design was focused on identifying within-model sex-specific and disease-stage dependent alterations relevant for ameliorating AD progression; inclusion of age-matched genetic background control C57BL/6 mice could enhance the interpretability and translational relevance of the findings. Second, we did not measure circulating gonadal hormones since our study design did not include repeated sampling or estrous cycle staging. A single-point hormone measurement would not have yielded biologically meaningful data due to the large physiological variability inherent in their sampling. Third, we only measured ectopic lipid deposition in the liver, so therefore it is possible for lipids to accumulate in other organs such as muscle. Finally, we focused on specific comparisons of sham and VFR surgery animals to their age- and sex-matched controls, consistent with the hypothesis, by means of a t-test. This was done to avoid exploratory-type findings for possible age and sex effects since our primary goal was to assess whether VFR improved cognitive performance at distinct disease stages within each sex.

